# Proneural lineage identity is the primary determinant of embryonic synchrony patterns in the developing hindbrain

**DOI:** 10.1101/2025.11.14.688499

**Authors:** Matthias Blanc, Lydvina Meister, Gonzalo Ortiz-Álvarez, William C. Lemon, Ulla-maj Fiuza, Philipp J. Keller, Isabel Espinosa-Medina, Cristina Pujades

## Abstract

Does the stem cell origin impact on how daughter neurons acquire functional characteristics? Here, we uncovered that progenitor identity, defined by proneural gene expression, hardwires the functional mode (synchrony vs. asynchrony) of a neuron. Multimodal cell lineage reconstruction combining intersectional lineage tracing, newly developed computational tools, and genetic knockouts revealed that different progenitors produced functionally distinct neuronal subtypes in the zebrafish hindbrain. Dynamics of neuronal production suggest that progenitor competence changes over time. Whereas *neurog1*-expressing progenitors contributed to both glutamatergic and GABAergic lineages at early embryonic stages, later, other progenitor pools also assumed this role. Whole-hindbrain 3D atlases, in which we combined calcium imaging to monitor spontaneous neuronal activity with genetic perturbations and progenitor origin, unveiled that the emergence of neuronal activity patterns was presaged by their progenitor identity. This suggests a link between cell ontogeny and neuronal activity in the zebrafish hindbrain, and more fundamentally how the embryonic progenitor identity controls the emergence of neuronal diversity in glutamatergic and GABAergic neuronal populations.

## INTRODUCTION

The survival of an organism requires the continuous coordination of the activities of its organs with the ever-changing external environment that it encounters. The responsibility for these tasks falls on the mature differentiated neurons, which are generated during embryonic development, and whose activity reflects the structural and functional organization of the emerging network^1,2^. While much is known about how neurons are generated^3^ and what neurons transmit different neurotransmitters, we still do not know why neurons with the same neurotransmitter have different activity modes.

During the development of the Central Nervous System (CNS), several neuronal types originate from more than one source following differential spatial cues^4,5^, being the spinal cord the most studied model^6,7^. However, studies of transcriptional trajectories and lineage analysis on the anterior mammalian brain suggest that there is not a common blueprint for how different neuronal subtypes emerge from multipotent neuronal precursors^8–10^. Temporal mechanisms also play a large role in controlling progenitor competence^11^. For instance, cortical progenitors mature over time such that we find inverted temporal gradients of “early” vs. “late” genes^12^, and neural progenitors gradually switch from specifying earlier-born cell fates to specifying later-born neurons^12–14^. In the hindbrain, the neuronal spatial distribution foreshadows the time of differentiation^15^, and specific neuronal functions have been correlated to neuronal birthdate^16,17^. Whereas previous brain studies have linked clonal relationships and neuronal activity^18^, or cell profiling and activity patterning^9^, we still do not know what underlies the emergence of specific neuronal activity patterns.

Addressing how this occurs has been technically challenging to date since it is difficult to trace the origin of functional neurons while recording their spontaneous activity. However, the zebrafish hindbrain offers excellent experimental suitability since its transparency and small size allow high-resolution live imaging and genetic lineage tracing at whole-organ scale, while genome editing enables precise perturbations. This makes it possible to integrate cell lineage and function *in vivo* at the organ level. Further, the hindbrain is highly conserved across vertebrate evolution^19^ and neurogenesis occurs extremely rapid at early embryonic stages^15,20^ establishing functional circuits essential for its fundamental physiological responses such as respiration, circulation, arousal and movement^21–23^.

Here, we present evidences to propose that proneural lineage identity is the primary determinant of embryonic synchrony patterns in the developing hindbrain. By combining progenitor intersectional fate mapping^24^ and CRISPR-based perturbation experiments with a new computational pipeline to generate 3D reference maps of whole brains, we establish the differential temporal contribution of distinct progenitor pools –defined by proneural gene expression– to the glutamatergic and GABAergic lineages. We find that both types of neurons arise from multiple proneural progenitor pools and that their production is not temporally synchronized. Further, by using 3D spatial correlation and calcium imaging to record neuronal activity in the entire developing hindbrain, we show that neuronal activity patterns are associated with progenitor identity. Finally, by performing calcium recordings upon perturbation of distinct progenitor pools we prove that neurons with distinct neuronal activity features originate from temporally and spatially defined progenitor populations: early-born, medially located progenitors produce neurons characterized by high-amplitude, low-frequency, and synchronized activity, consistent with a role in establishing the baseline neuronal activity; by contrast, later-born, laterally located progenitors produce neurons with low-amplitude, high-frequency, and asynchronous activity, suggesting contributions to later functions. Thus, by using this multimodal approach, we unveiled that the progenitor identity is the primary determinant of the spontaneous activity mode (synchrony vs. asynchrony) of a neuron in the developing hindbrain, supporting the idea that the lineage and developmental trajectory of a cell are key contributors to their neuronal activity.

## RESULTS

To reveal how spatial neurogenesis patterns are established in the hindbrain progenitors, we first assessed the temporal expression of proneural genes (PN) –transcription factors that commit progenitors towards the neuronal lineage. At 24hpf, PN expression differed along the DV axis (Figure 1a–e), with 5 distinct progenitor pools defined according to the PN-gene combinations. While most progenitor pools’ distribution was similar to the spinal cord^25,26^, the most dorsal *neurog1* expression domain was absent (Figure 1a, e). *neurog1*-expressing progenitors were mainly, but not exclusively, contained within the *ascl1b*-domain (Figure 1a–b, e), whereas *ptf1a* and *atoh1a* progenitors were segregated to the dorsal-most part of the tissue with no observable overlap with other PN genes (Figure 1c–e; ^27^). Progenitors placed in the dorsal domain at 24hpf were located dorsolateral at 48hpf due to the progressive opening of the neural tube (Figure 1f–j). By 48hpf, most of the *neurog1*-domain was still contained within the *ascl1b*-region (Figure 1f–g, j), which expanded laterally towards the *ptf1a*-domain (Figure 1g–h, j). This change in the temporal PN expression profile may result in part from two intertwined morphogenetic events that occur during development, the growth of the neuronal differentiation domain at the expense of the progenitor territory^15,28^ and the formation of the brain lumen^27–29^. The expression of PN genes differed along the anteroposterior (AP) axis (Figure S1a–j). At 24hpf, *ascl1b* was expressed all along the AP axis in a ventral domain comprising *neurog1* (see dorsal views in Figure S1a–b, e; a’–b’, e’) and only in the posterior hindbrain this expression was comprised in a specific dorsal domain in between the *ptf1a* and *atoh1a* (Figure 1b, e). By 48hpf, this dorsal *ascl1b* domain displayed *neurog1* as well, while still being absent in more anterior regions (Figure 1f–g, j; Figure S1f–f’, j–j’). Overall, these results show that PN genes dynamically pattern the neural progenitor domain (Figure 1k).

**Figure 1:**
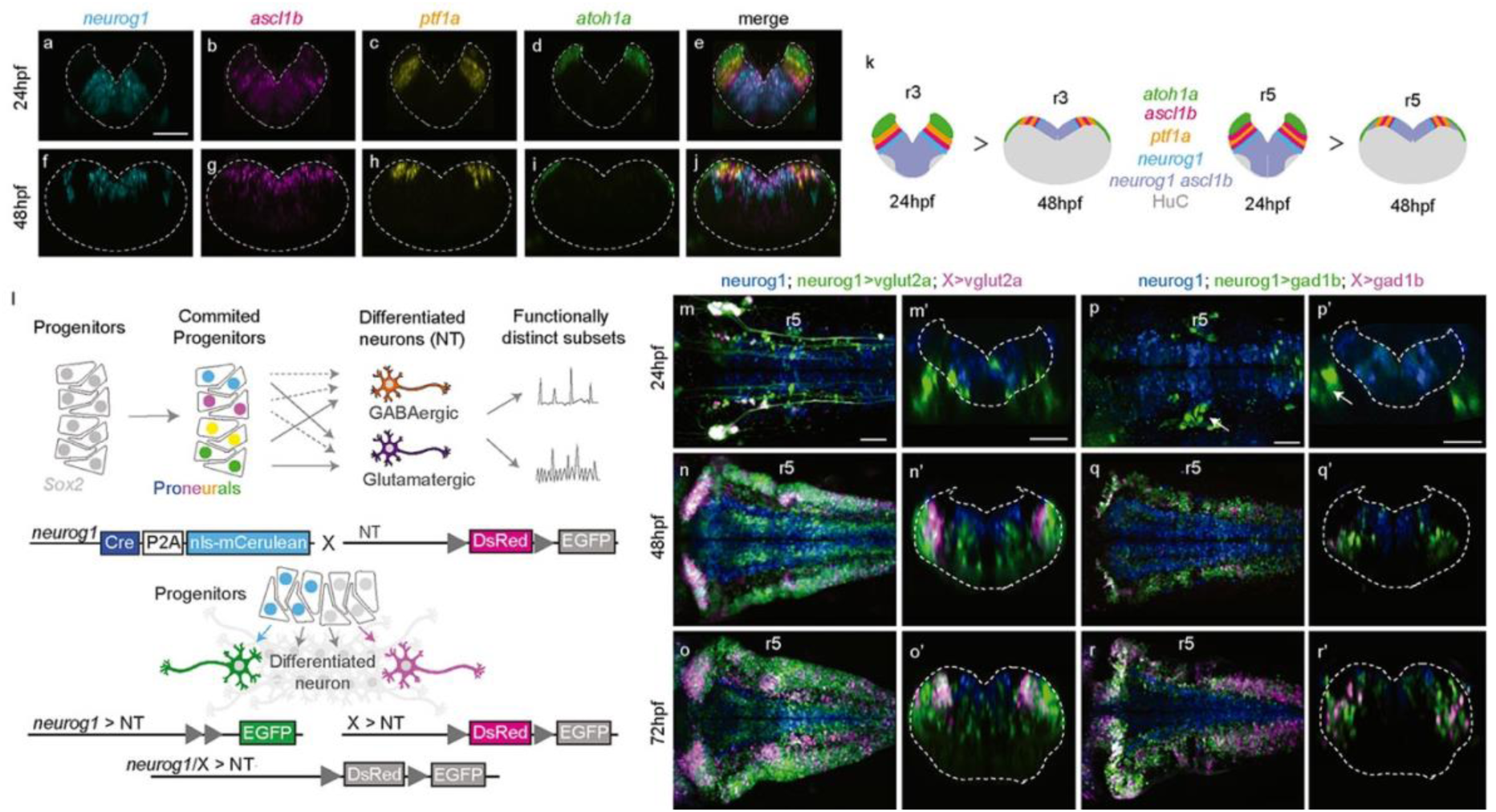
*neurog1*-progenitors differentially contribute to glutamatergic and GABAergic lineages. a–j, Multiplex HCR of *neurog1*, *ascl1b*, *ptf1a* and *atoh1a* proneural genes at 24hpf (a–e) and 48hpf (f–j). Transverse views of single channels (a–d, f–i), and the merge of all channels (e, j). Note the differential expression of proneural genes along the dorsoventral (DV) axis at 24hpf and mediolateral (ML) axis at 48hpf. Transverse views through rhombomere (r) 5. k, Scheme summarizing the differential expression of proneural genes along the DV and ML axes of the hindbrain together with the neuronal differentiation domain (HuC) along rhombomeres 3 and 5. l, Scheme depicting the goal of the work, dissecting the contribution of different progenitor pools to excitatory (glutamatergic) and inhibitory (GABAergic) neurons, and the employed strategy to address the contribution of *neurog1*-progenitors to both neuronal lineages. m–r, m’–r’, Production of glutamatergic (m–o) and GABAergic (p–r) neurons over time, according to the progenitor origin (*neurog1*-derivatives in green, *non-neurog1*-derivatives in magenta) at the indicated developmental stages. Dorsal views with anterior to the left (m–r) and transverse views through rhombomere 5 (m’–r’). White arrows in (p, p’) indicate the statoacoustic sensory ganglion. hpf, hours post-fertilization; NT, neurotransmitter; r, rhombomere; X, unknown PN-expressing progenitors.

Next, to monitor the growth of the *neurog1*-progenitor pool we built up the zebrafish Tg[neurog1:Cre-P2a-nlsCerulean] transgenic line, with Cre under the control of *neurog1* promoter. In transgenic embryos, *neurog1* mRNA expression was restricted to the progenitor domain (Figure S2a–c, a’–c’), and accordingly, HCR analysis revealed overlapping expression of *Cre* and *neurog1* (Figure S2d–d’’, e–e’’). *In vivo* Light Sheet Fluorescence Microscopy (LSFM) imaging of Cerulean expression demonstrated this new transgenic line faithfully recapitulates the expected neurog1 spatiotemporal patterning during hindbrain morphogenesis (Figure S2f–j, f’–j’).

### *neurog1*-progenitors contribute to both glutamatergic and GABAergic neurons with distinct temporal dynamics

We explored the differential temporal contribution of distinct PN-expressing progenitor pools to the excitatory (glutamatergic) and inhibitory (GABAergic) neuronal populations, which constitute 80% of the overall hindbrain neurons born before 72hpf^15^. We used progenitor-restricted intersectional mapping^24^ to singularize cell lineages arising from specific progenitor populations and assessed their temporal contribution to the differentiated neuronal derivatives (Figure 1l).

To address the contribution of *neurog1*-progenitors to the specific glutamatergic (vglut2a) and GABAergic (gad1b) neuronal subsets, Tg[neurog1:Cre-P2a-nlsCerulean] fish (hereafter Tg[neurog1:Cre]) were crossed with either Tg[vglut2a:Lox-DsRed-Lox-GFP] or Tg[gad1b:Lox-DsRed-Lox-GFP] fish^30^ (hereafter Tg[vglut2a:switch] and Tg[gad1b:switch], respectively) and the resulting embryos were *in vivo* imaged during periods encompassing the beginning of neuronal differentiation (24hpf) until most of the hindbrain cells are postmitotic neurons (72hpf)^15,28^. The production of glutamatergic and GABAergic neurons followed distinct dynamics with several progenitor pools contributing to their growth (Figure 1m–o, p–r; see *neurog1*-derived cells in green and the ones derived from other progenitors in magenta). *neurog1*-progenitors generated both types of differentiated neurons; whereas glutamatergic derivatives were already widely present in the hindbrain by 24hpf (see green cells in Figure 1m, m’) no GABAergic neurons were there (Figure 1p, p’). The only GABAergic neurons present at this stage derived from *neurog1*-cells and corresponded to sensory ganglia adjacent to the hindbrain^31,32^ (see white arrow in Figure 1p, p’). By 48hpf, although *neurog1*-progenitors were still generating glutamatergic neurons, other PN-expressing progenitors started producing them (see magenta cells in Figure 1n, n’) and this was maintained by 72hpf (Figure 1o, o’). On the other hand, by 48hpf most of GABAergic neurons originated from *neurog1*-progenitors (see green cells in Figure 1q, q’), and by 72hpf, GABAergic neurons were mostly generated by other progenitor pools (see magenta cells in Figure 1r, r’). To further address the progenitor contribution dynamics, we performed LSFM recordings and observed two distinct behaviors for these neurotransmitter’s defined subsets. Glutamatergic neurons were simultaneously and consistently produced by both *neurog1*-expressing progenitors and other pools across development (Videos 1–2), whereas GABAergic neurons were exclusively produced by *neurog1*-progenitors early on, and later by other contributors (Videos 3–4). Although the secondary production of GABAergic neurons was initiated later, it resulted in the production of a relatively similar proportion of neurons (Videos 3–4). These results demonstrate a differential temporal growth of glutamatergic and GABAergic neurons, with distinct spatiotemporal contribution of progenitor pools. Whereas *neurog1*-progenitors contribute to both lineages in early embryonic stages, later, other progenitor pools either also assume this role (for excitatory neurons), or take over (for inhibitory neurons). Next, we wanted to identify the other progenitor pools contributing to these lineages.

### Glutamatergic and GABAergic lineages derive from distinct progenitor pools’ combinations

To resolve the heterogeneity of neuronal contributors we combined progenitor-restricted intersectional mapping using the Tg[neurog1:Cre] and the Tg[vglut2a:switch] and Tg[gad1b:switch] lines with a CRISPR/Cas9-based approach that produces loss-of-function (LOF) phenotypes in F_0_ by redundantly targeting single genes^33,34^ –and therefore impair the formation of specific progenitor pools. Either *neurog1*, *ascl1b*, *ptf1a* or *atoh1a* proneural genes were targeted by injecting one cell stage embryos with four different gRNAs per proneural gene (Figure S3a–c), and we assessed the impact on i) the expression of PN genes in the adjacent domains by HCR-multiplex transcript analysis (Figure 2a; Figure S3d–m), ii) the glutamatergic and GABAergic lineages (Figure 2b), and iii) the cell death, showing negligeable apoptosis associated to CRISPR-effects in the neural tube (Figure S4). *neurog1*_gRNAs injections resulted in smaller hindbrains with no dramatic changes in adjacent PN gene expression at 24hpf (Figure 2a; Figure S3d–e, i–j), and an upregulation of *neurog1* in the statoacoustic ganglion as previously reported^31,32^. *ascl1b*_gRNAs injected embryos displayed smaller hindbrains with a smaller *neurog1* territory at 24hpf (Figure 2a; Figure S3d, f, i, k) suggesting a regulation of *neurog1* by *ascl1b*. *ptf1a*_gRNAs resulted in an expansion of *ascl1b* (Figure 2a; Figure S3d, g, i, l) indicating that *ptf1a*- and *ascl1b*-domains interact. Finally, *atoh1a*_gRNAs did not dramatically impact the adjacent *ptf1a* domain (Figure 2a; Figure S3d, h, i, m) showing that there was no cross-regulation as previously demonstrated in the *atoh1a^fh282^* mutants^27^.

**Figure 2:**
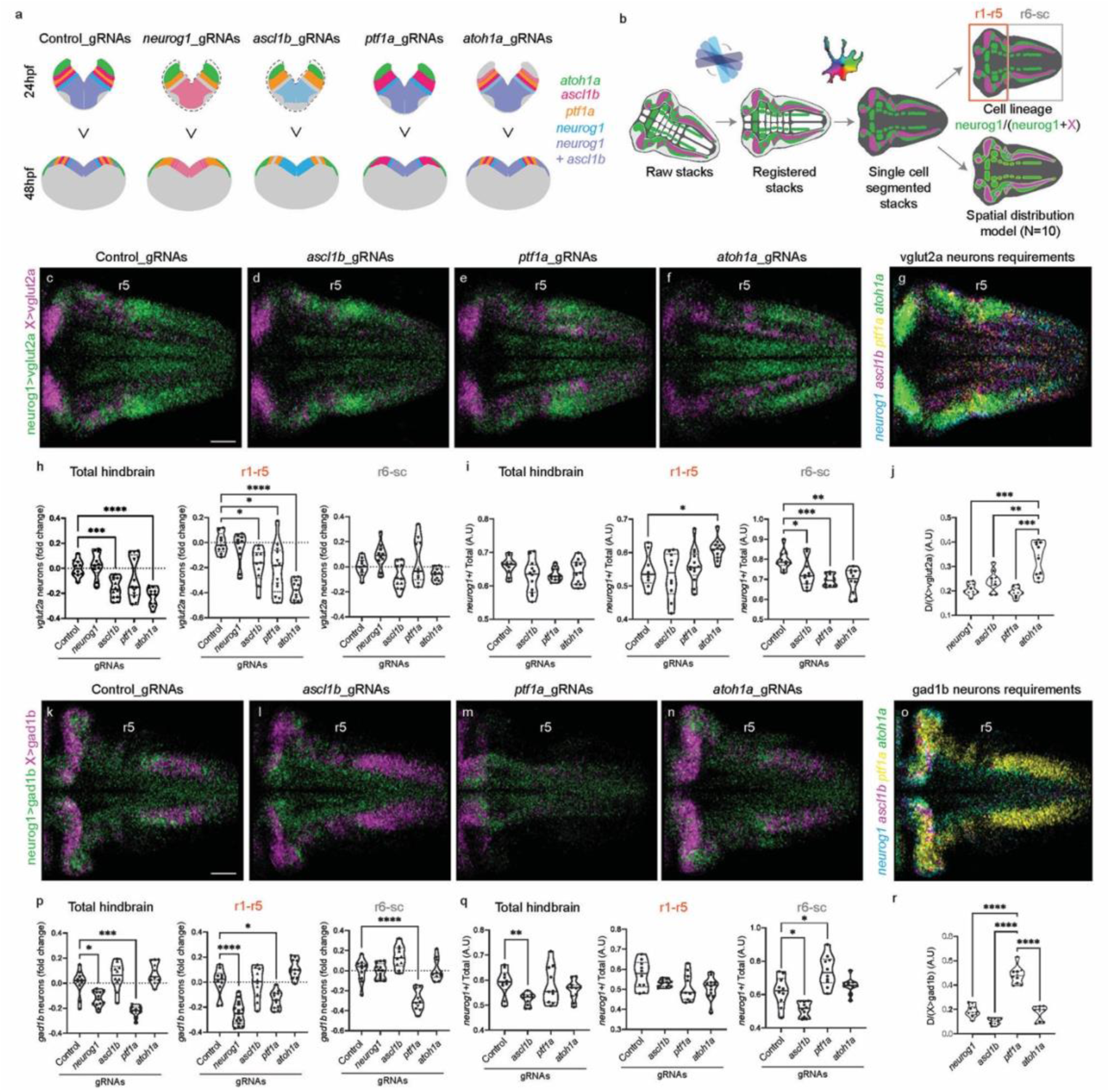
Glutamatergic and GABAergic lineages derive from distinct progenitor pools’ combinations. a, Summary of the observed phenotypes by multiplex HCR upon proneural gene disruption (Figure S3). b, Scheme depicting the processing of the images. Hindbrains from embryos injected with the corresponding gRNAs were imaged at 72hpf and images from 10 hindbrains per condition underwent alignment. Green (*neurog1*-derived) and magenta (*non neurog1*-derived) cells were segmented using Cellpose to i) quantify the proportions of glutamatergic and GABAergic neurons derived from the different progenitor populations and ii) construct spatial distribution 3D models allowing the evaluation of the impact of LOF on the neuronal populations across the tissue. c–f, Spatial distribution 3D-models of glutamatergic cells derived from *neurog1-*(green) and *non-neurog1* progenitors (magenta) upon the indicated conditions. g, 3D-map showing the progenitors’ origin requirements for glutamatergic neurons. Note that the main contributors are n*eurog1*-, a*scl1b*-and a*toh1a*-cells. h–i, Violin plot showing the variation on the proportion of all glutamatergic neurons and *neurog1*-derived glutamatergic neurons, respectively, upon PN_gRNAs injections with reference to the control, in the overall hindbrain (total hindbrain), and specifically in the anterior hindbrain (r1-5) or in the posterior hindbrain (r6-sc). j, Requirement of the *non-neurog1* derived glutamatergic neurons for each of the proneural progenitor pools, measured through the intersection of control samples with the lineage requirements maps and displayed as proportion. k–n, Spatial distribution 3D-models of GABAergic cells derived from *neurog1*-(green) or *non-neurog1* progenitors (magenta) upon the indicated conditions. o, 3D-map showing the progenitors’ origin requirements for GABAergic neurons. Note that the main contributors are n*eurog1*-, *ptf1a*-cells. p–q, Violin plot showing the variation on the proportion of all GABAergic neurons and *neurog1*-derived GABAergic neurons, respectively, upon PN_gRNAs injections in reference to the control, in the overall hindbrain (total hindbrain), and specifically in the anterior hindbrain (r1-5) or the posterior hindbrain (r6-sc). r, Requirement of the *non-neurog1* derived GABAergic neurons for each of the proneurals’ progenitor populations, measured through the intersection of control samples with the lineage requirements maps and displayed as proportion. For individual sample measurements Brown-Forsythe and Welch one-way Anova tests were performed to account for variations of standard deviation and followed by Dunnett’s multiple comparisons test (h–i; p–q; Supplementary Dataset 1). For the requirement maps intersections to individual lineage traced samples, a repeated measurement (RM) one-way Anova test with Geisser-Greenhouse correction was performed to account for matched dataset followed by a Holm-Sidak’s multiple comparison test (j, r; Supplementary Dataset 1). All images display dorsal views of the hindbrain with anterior to the left. r, rhombomere. *p<0.0332; **p<0.0021; ***p<0.0002; ****p<0.0001

Next, we analyzed the impact of proneural gene LOF on the glutamatergic and GABAergic lineages by 72hpf (Figure 2b). Double transgenic embryos, Tg[neurog1:Cre:vglut2a:switch] or Tg[neurog1:Cre:gad1b:switch], were injected with control or PN_gRNAs, imaged at 72hpf, and cell lineages’ analyses were performed to i) quantify the proportions of glutamatergic and GABAergic neurons derived from the different progenitor populations and ii) construct spatial distribution models allowing the evaluation of the impact of LOF on the neuronal populations across the tissue (Figure 2b). To quantify the outcome on the neuronal populations we established a computational workflow combining i) a newly generated deep learning-based software, RAD (Rigid Alignment with Deep learning) to perform rigid registration among samples, which enabled supervised or unsupervised alignment of tissues using a trained digital neural network (see https://github.com/cristinapujades/Blanc_et_al_2025), ii) the latest version of Cellpose^35^, a deep learning single cell segmentation framework, and iii) a set of Python scripts to quantify, process and generate representative 3D-models averaging 10 hindbrains per condition including different transgenic backgrounds (Figure 2b). Our averaged 3D-models revealed an important decrease of glutamatergic neurons upon *atoh1a* downregulation (Figure 2c–f). Quantification of individual samples showed that the global glutamatergic population was significantly reduced upon *ascl1b* and *atoh1a* LOF and that this reduction was mostly due to a cell population located in the anterior hindbrain (r1-r5) (Figure 2h). A minor reduction was also observed anteriorly upon *ptf1a* LOF, demonstrating an indirect effect of the GABAergic perturbation (Figure 2h). The shift in the proportion of *neurog1*-derivatives in the r1-r5 region placed *atoh1a*-progenitors as the major other contributor to the glutamatergic pool (Figure 2i). To spatially map these progenitors, we put forward the depleted and compensated cell subsets across the hindbrain by performing deltas of spatial distribution between control and LOF 3D-models (Figure S5a–d), and thus generated a 3D global model of PN-progenitor requirements for glutamatergic neurons across the hindbrain, which informed us about the spatial requirements for *neurog1*-, *ascl1b*-and *atoh1a*-progenitors within the final glutamatergic neuronal population (Figure 2g). These results indicate that the several progenitor pools that generate glutamatergic neurons are not segregated progenitor subsets, except for *atoh1a*-progenitors, indispensable to the dorsolateral neurons (Figure 2g). When *non-neurog1* derivatives were overlapped with the proneural gene requirements’ map, the quantifications revealed that this population was significantly enriched in the *atoh1a*-dependent subset, bringing further confirmation that *atoh1a* was the other major progenitor pool producing glutamatergic neurons (Figure 2j).

When the same strategy was applied to the GABAergic neurons, we observed a dramatic decrease of this neuronal population upon downregulation of *ptf1a* (Figure 2k–n). Quantifications revealed a significant reduction of GABA neurons upon *neurog1* and *ptf1a* LOF (Figure 2p), which was restricted to the anterior hindbrain in the case of *neurog1*_gRNAs but was evenly distributed along the AP axis in the *ptf1a*_gRNAs (Figure S5e–h). To further confirm that the *neurog1*-population was able to produce GABAergic neurons independently of *ptf1a*, we performed LSFM imaging in the cell lineage tracing reporter background and observed that indeed *neurog1*-cells were giving rise to GABAergic neurons in the absence of *ptf1a* (Video 5). Moreover, we observed a decrease in the ratio of *neurog1*-derived neurons upon *ascl1b* LOF in the whole hindbrain with no significant impact on the global GABAergic cell quantity (Figure 2q) unveiling a compensation mechanism and suggesting that at least a subset of the *neurog1*-progenitors depended on *ascl1b*. When the progenitors’ origin requirement maps were confronted to our lineage tracing models, we further confirmed *ptf1a* to be the most likely major other contributor to the GABAergic lineage (Figure 2o, r). These results back up our previous observation that *neurog1*-progenitors are the main producers of GABAergic neurons at early stages, and that later, *ptf1a*-progenitors take up that role. Accordingly, the *neurog1*-expression domain located laterally, between the *atoh1a* and the *ptf1a* domains, disappeared upon *ascl1b* or *ptf1a* LOF, as shown by HCR experiments at 48hpf (see asterisk in HCR for *neurog1* in Figure S3i, k–l). Overall, these observations suggest that the GABAergic neurons originating from *neurog1*-progenitors derive from two subpopulations, one located medially that is *ptf1a*-independent and one located laterally that is *ptf1a*- and *ascl1b*-dependent.

### Progenitor identity defines the early activity patterning of hindbrain neurons

To understand the extent to which the ontogeny of a cell is linked to its function, we first addressed the activity of the cell populations in the embryonic hindbrain at 72hpf. Since at this stage no neuronal activity recordings were publicly available, we made use of the Tg[elavl3:jrGECO1b] transgenic line to record spontaneous neuronal activity in the whole hindbrain (Figure 3a; Figure S6a) using high-speed simultaneous multi-view light-sheet microscopy (hs-SiMView^36^). To extract single cell information, we developed an image analysis pipeline to establish individual cell profiles of the detectable active neurons across the whole hindbrain (Figure 3a; Figure S6b). Using Louvain clustering we detected 3-4 communities (C1– C4) per sample and identified 3 conserved activity patterns across all analyzed samples with matching features that we called matching communities (Figure 3b–c, M1–M3; see M1 in red, M2 in green and M3 in blue in Figure 3c). Each matching community displayed a different synchronization score, matching community M3 being the least synchronized (Figure 3d; M1 = 0.33, M2 = 0.21, M3 = 0.06). Moreover, matching communities plotted consistently in a segregated manner upon UMAP analysis (Figure 3d). To explore the position of neurons of the matched communities, we plotted their spatial distribution and revealed that indeed neuronal activity features were specifically allocated within the hindbrain (Figure 3e–f). The matching community M3, located in the most lateral domain (compare Figure 3f with Figure S6c–e; sites of *ptf1a*- and *atoh1a*-dependent neurons) displayed very low coherence (i.e. synchronicity within its own community, Figure 3g). Dissecting the neuronal composition of these matching communities revealed that most of the active neurons in M1 to M3 were glutamatergic and GABAergic neurons, although there was a small proportion of unidentified neurons (Figure 3h, see grey bars). Nevertheless, when the lineage contribution to glutamatergic and GABAergic populations was analyzed noticeable differences in community configuration depending on the progenitor origin were observed, namely more *neurog1*-derived cells in M1 and M2 (Figure 3h; M1 82% vs M2 80% vs M3 69%). More specifically, M3 was enriched in *ptf1a*-dependent GABAergic neurons and *atoh1a*-dependent glutamatergic neurons (Figure 3i, see in M3 *ptf1a>gad1* and *atoh1a>vglut2a*). When we analyzed distinct neuronal activity features according to lineage contribution, we observed that *neurog1*-derivatives showed a higher average event amplitude and synchronicity score (Figure 3j–k). Markedly, *ascl1b*-dependent glutamatergic neurons displayed higher synchronicity than the ones requiring *atoh1a* (Figure 3l). Finally, we wanted to identify the leading cause (neurotransmitter, lineage or birthdate) of the changes in emergent activity features. For this, we computed the shapely share of activity features (see Matherials and Methods), and lineage emerged as the dominant source of variability for event amplitude and synchronicity (Figure 3m–n). Thus, these correlations suggest that lineage is the main responsible of the synchronicity feature in the early embryo.

**Figure 3:**
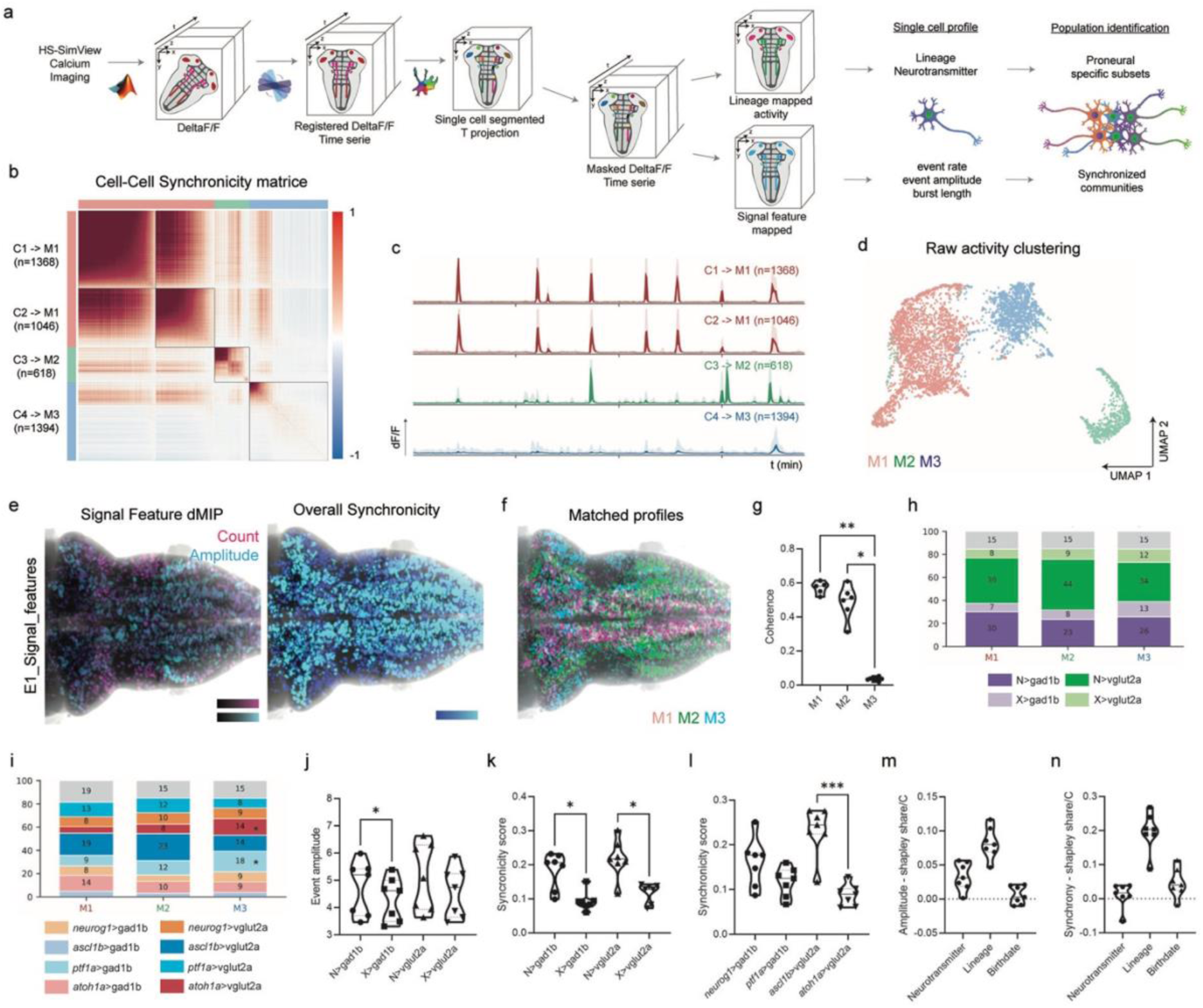
Activity neuronal features rely on the progenitor origin. a, hs-SimView recordings are first processed through a series of Matlab scripts to extract the deltaF/F, and the subsequent recording is then aligned to our reference atlas using our alignment software. Aligned recordings of deltaF/F are projected over time to obtain a 3D image representing neuron maximal activity over the time sequence. The projection is passed to Cellpose for single cell segmentation and intersected with our previously established lineage maps. Each cell’s deltaF/F signal variations over the time sequence are analyzed to obtain feature maps reflecting the cells’ activity characteristics. b, Cell-cell synchronicity matrices of hindbrain active neurons. Note that we can identify 4 different neuronal communities (C1–C4) matched to 3 conserved activity profiles called matching communities (M1–M3). c, Neuronal activity profiles of C1–C4 indicating the number of segmented cells constituting each community (n) and the activity profile they were matched (M1–M3). d, UMAP displaying the 3 different populations according to their activity profiles (M1–M3) within a representative sample. e, Representative intensity projections of 3D-maps of active neurons according to different features: count and amplitude or overall synchronicity. f, Intensity projections within the embryonic hindbrain of the 3D-map of the activity profiles distribution, color-coded according to M1–M3. g, Violin plot showing the coherence, or within group synchronicity, of cell populations displaying a given matched activity profile. h, Box plot displaying the cell lineage composition of M1–M3. i, Box plot displaying the progenitor origin requirements of M1–M3. j–k, Violin plots showing the event amplitude and synchronicity score observed according to the progenitor origin mapping. l, Violin plot showing the observed synchronicity score according to the progenitor origin requirements. m–n, shapely share/C of each map for synchrony and event amplitude. Values below 0 mean that adding the map reduces out-of-sample prediction. For activity features per lineage measurements, non-parametric Friedman tests were performed and followed by Dunnett’s multiple comparisons test (j–k). *p<0.0332; **p< 0.0021; ***p<0.0002; ****p<0.0001.

To demonstrate a causal relationship between the presence of cell lineages and the activity patterns observed across the hindbrain, we combined calcium imaging recordings with the CRISPR/Cas9-based loss-of-function approach. We observed major changes in neuronal activity (Videos 6–10) such as a decrease in cell-cell activity correlations (i.e synchronicity) upon *ascl1b* loss of function (Figure 4a–a’, c–c’). This was further confirmed by the analysis of the average activity pattern over communities (Figure 4a’’–e’’), with *ascl1b*_gRNAs embryos showing a lack of temporally defined spiking and no synchronicity (Figure 4a’–a’’, 4c’–c’’). *neurog1*_gRNAs embryos displayed an overall decrease in amplitude but no loss of synchronicity (Figure 4a’–4b’). Neuronal activity upon *ptf1a* loss-of-function resulted in wider spikes without losing tissue-wide synchronicity or amplitude of signal (Figure 4a’’, 4d’’), with the possibility that the generation of spiking trails could be associated to the direct disruption of spike timings resulting from an imbalanced inhibition^37^. On the contrary, upon *atoh1a* loss-of-function no loss of synchronicity or amplitude of signal were observed, whereas the number of spikes was greatly increased across the whole tissue, although not in the form of spiking trails but as individual events (Figure 4a’’, 4e’’). We further observed that upon LOF of *ascl1b*, cells felt into the M3 category (Figure 4c’’, f), and upon quantification we verified that LOF of *ascl1b* resulted in drastic decrease of both synchronicity score and average event amplitude (Figure 4g–h). Since a*scl1b* and *neurog1* were expressed in similar domains (Figure 1; Figure S3), we quantified the synchronicity score in *neurog1*-derived populations within the *ascl1b* LOF and observed a significant decrease (Figure 4i), supporting the hypotheses that the high synchronicity score of *neurog1*-derivatives (Figure 3k) was mainly due to their overlap with the *ascl1b*-progenitors, and that *ascl1b* progenitors drived the emergence of the synchronized spontaneous activity in the early embryo hindbrain.

**Figure 4:**
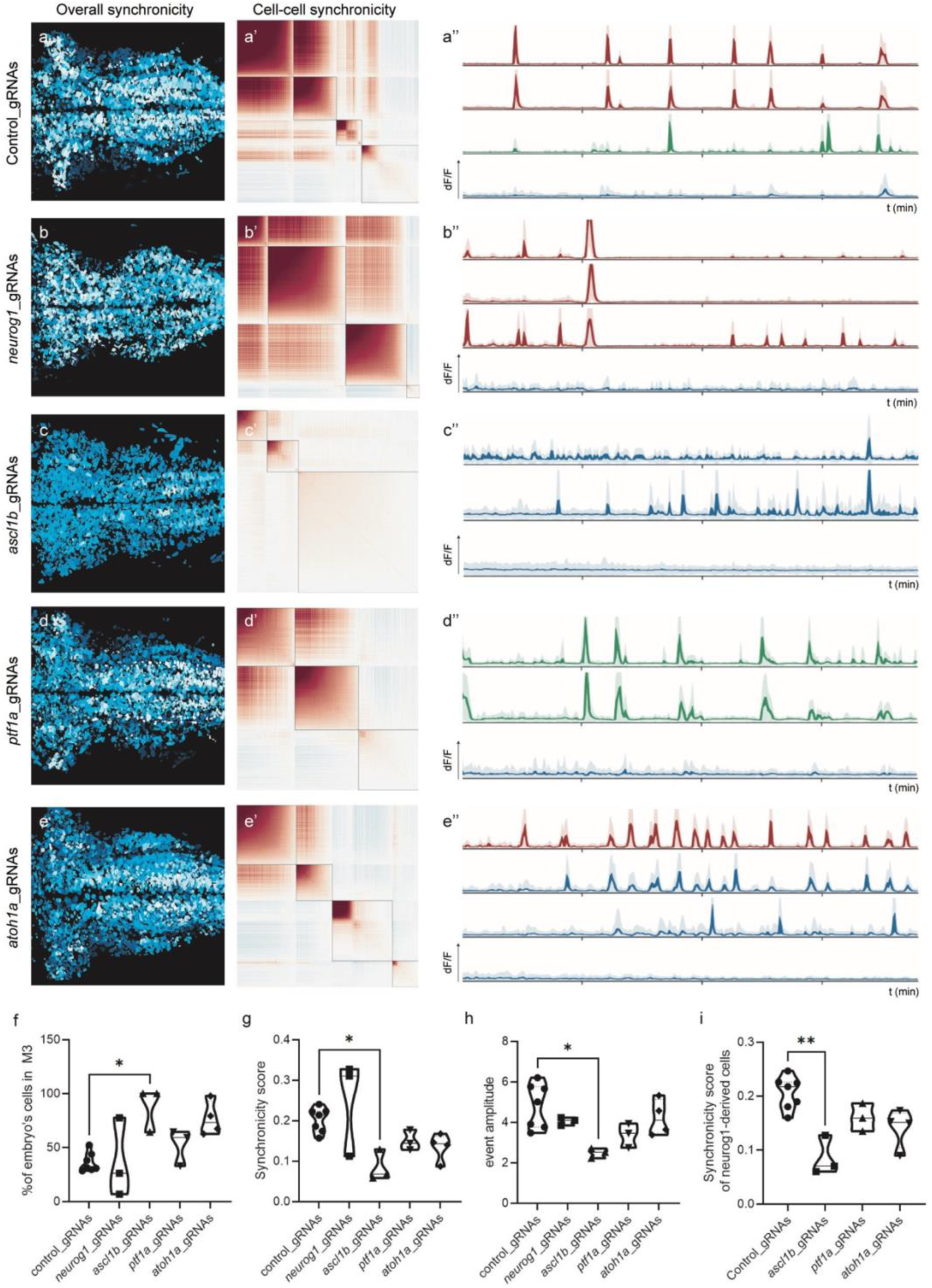
Disruption of specific progenitor pools results in differential changes in neuronal activity. a–e, Synchronicity of neuronal activity in control hindbrains and upon disruption of proneuronal progenitor pools (Videos 6–10). Dorsal views with anterior to the left. a’–e’, Cell-cell synchronicity maps showing the Louvain communities upon different conditions. a’’–e’’, Hindbrain neuronal activity profiles of each community and their corresponding matching to the previously described conserved activity patterns or matching communities (M1–M3) under the different conditions. Note how disruptions in the progenitor origin result in changes in community distribution and neuronal activity. f, Violin plot of the percentage of each embryo’s cells displaying the M3 activity profile. g–h, Violin plots showing the event amplitude and synchrony scores observed in each embryos i, Violin plot of the synchronicity scores within the *neurog1*-derived mapped subset.

## DISCUSSION

Neuronal identity is defined by specific gene expression programs, which are executed by gene regulatory networks^38^. Further, the cell lineage history together with a specific spatiotemporal trajectory within complex tissues are emerging as central to defining cell fate1139. Here, using a new multi-modal 3D strategy in the zebrafish hindbrain, we provide evidence to respond to the question why neurons with the same neurotransmitter have different activity modes. We propose that the answer mostly lies in lineage and not in the neurotransmitter type. We find that glutamatergic and GABAergic neurons derive from different proneural progenitor pools and we hypothesize that this proneural lineage identity is the primary driver of the emergence of functional neuronal diversity (Figure 5), providing *in vivo* evidence for a link between neuronal cell ontogeny and activity.

**Figure 5:**
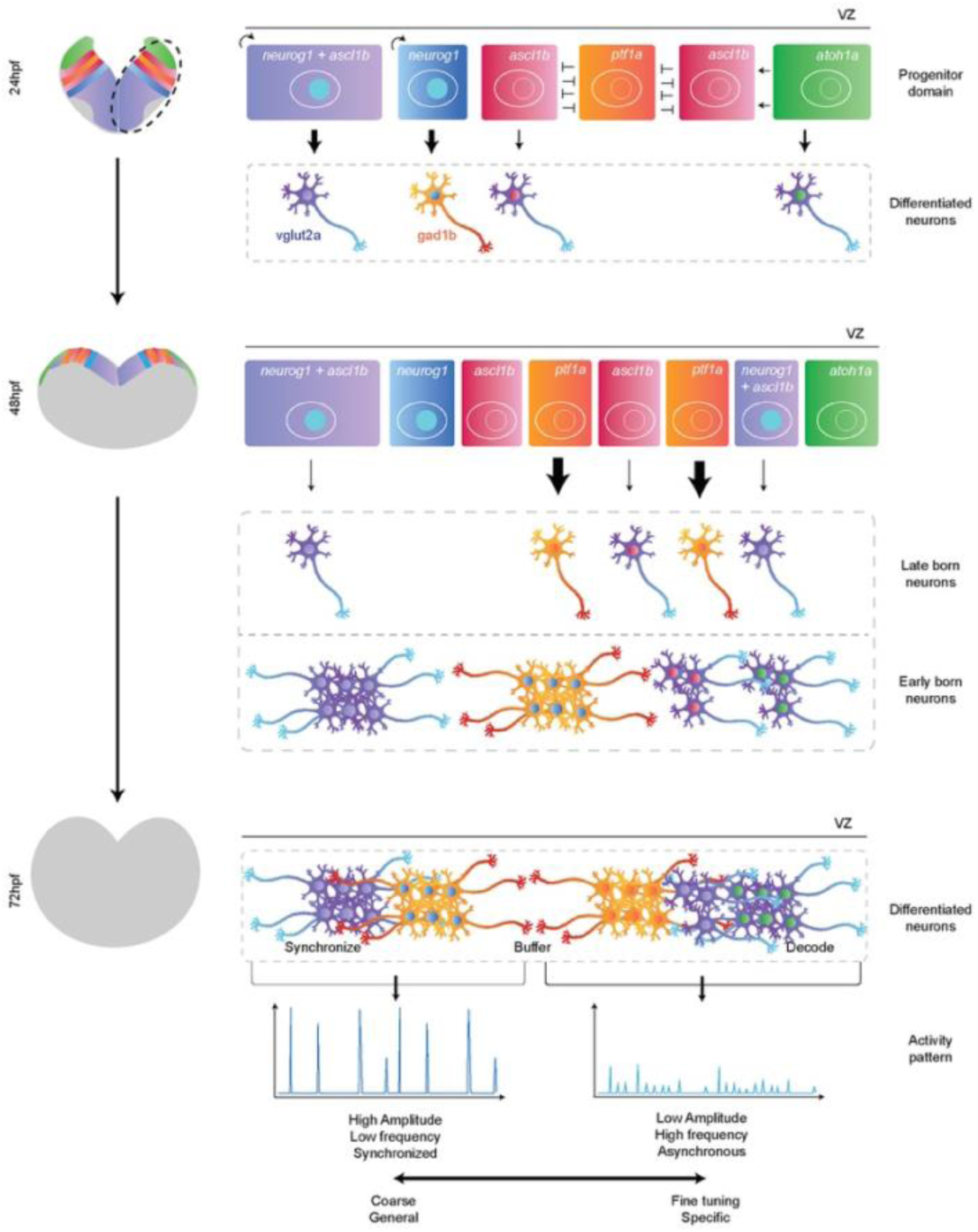
Progenitor spatiotemporal identity is crucial for establishing terminal characteristics such neuronal activity and neurotransmitter identity. The neuronal progenitor domain is patterned along the DV and ML axes by differential proneural gene expression. Early on (24hpf), *neurog1*-stem cells are the first one to produce glutamatergic (vglut2a) and GABAergic (gad1b) neurons with differential temporal dynamics. This is followed by *atoh1a*- and *ascl1b*-progenitors that are continuously added to the glutamatergic population, whereas *ptf1a*-progenitors take the relay for GABAergic neurons (48hpf). Our hypothesis is that this temporally defined progenitor identity determines specific neuronal activity features: early-born medially located progenitors produce neurons characterized by high-amplitude, low-frequency, and synchronized activity, likely establishing the baseline architecture; by contrast, later-born laterally located progenitors produce neurons with low-amplitude, high-frequency, and asynchronous activity, suggesting a role in more specific fine-tuning. VZ, ventricular zone; hpf, hours post-fertilization.

The production of glutamatergic neurons and GABAergic neurons does not follow a common rule in distinct areas of the CNS. In most anterior brain areas, there is a binary explanation of circuit function, with distinct neuronal types deriving from strictly segregated progenitor domains that express either *neurog1* (glutamatergic neurons) or *ptf1* (GABAergic neurons)^40,41^, whereas dorsal regions of the midbrain and the cerebellum contain intermingled populations of those two neuronal types^8^. We demonstrate that in the hindbrain both types of neurons derive from multiple progenitor pools with different temporal dynamics. Glutamatergic neurons derive mainly from *neurog1*, *ascl1b* and *atoh1a* progenitors whereas GABAergic neurons derive from *neurog1* and *ptf1a* progenitors, suggesting the presence of lineage-associated subtypes within those populations, as shown in other developmental systems such as the lymphatic vessels that derive from distinct groups of progenitors^42^. This indicates a certain developmental robustness of the progenitor pools during development, with limited compensation that nevertheless preserves lineage-associated differences. We further elucidate a crucial temporal component: at early embryonic stages, *neurog1*-expressing progenitors contribute to both lineages; in later stages, other progenitor pools either also assume this role (for glutamatergic neurons) or take over (for GABAergic neurons). Prior work suggested that neuronal age-related patterning provides a foundation for the assembly of hindbrain networks underlying diverse behaviors^15–17^. In this study, we provide complementary information about how progenitor identity, space, and developmental time jointly shape emerging patterns of neuronal activity in the hindbrain, supporting the notion that cell lineage and developmental trajectory are key contributors to cellular functional diversity. Further, our *in vivo* models integrate the temporal logic within intact tissue architecture helping us to better interrogate brain development while providing a foundation for future mechanistic studies.

Previous clonal studies on the mammalian cortex showed that the establishment of specific synaptic connections relied on the cell progenitor division mode and cell clonality^18^, highlighting the importance of the progenitor origin in setting a ground plan for neuronal circuits. Our evidence at the whole tissue scale is that the cell lineage and the spatiotemporal progenitor identity is essential to the emergence of patterned neuronal activity. The high reproducibility of neuronal activity patterns indicates the robustness of zebrafish embryo fundamental circuits for survival, in line with other studies showing that functional maturation of neurons is stereotyped, based on birth time and anatomical origin^43^. This does not mean that early acquired neuronal activity features will be maintained up to adulthood but rather that neuronal activities crucial for life-sustaining functions as those coordinated by the hindbrain are established very early. One of these examples would be the hindbrain V2a neurons, which are necessary for spontaneously occurring swimming as early as 3dpf and display region-specific differences in their ability to elicit or stop swimming behaviors^44^.

Therefore, we propose a model in which progenitor identity and lineage bias the activity of neuronal populations, more so than the neurotransmitter identity or birthdate of the neuron. This would be synchronizing for the most medial cells as shown upon disruption of *ascl1b*-progenitors, compensating for the neurons in between since loss of *ptf1a* results in reduced spike frequency and GABAergic neurons control action potential firing^37,45^, and decoding for cells in the most lateral domain receiving the sensory inputs. This suggests that the spatiotemporal patterning of the progenitor domain establishes neuronal diversity and constrains early neuronal activity, allowing the essential hindbrain resilience and adaptation necessary for its life sustaining functions. The combination of 3D-and *in vivo* experiments with new computational resources allowed us to build a common framework to globally interrogate the system and address previously inaccessible questions. In the future, combining this 3D lineage and activity map strategy with spatial transcriptomics or proteomics could further unveil the molecular mechanisms underlying the formation of specific neuronal activity traits.

## MATERIALS AND METHODS

### Ethics declarations and approval for animal experiments

All procedures were approved by the institutional animal care, the PRBB Ethics Committee in Animal Experimentation and the Departament de Territori i Sostenibilitat (Generalitat of Catalonia) in compliance with the National and European regulations. The PRBB animal facility has the AAALAC International approval B9900073. All the members accessing the animal house must hold the international FELASA accreditation. The Project License covering the proposed work (Ref 10642, GC) pays particular attention to the 3Rs. For the zebrafish work carried out at Janelia Research Campus (HHMI), all experiments were conducted according to the National Institutes of Health (NIH) guidelines for animal research and approved by the Institutional Animal Care and Use Committee of Janelia Research Campus (IACUC).

### Zebrafish strains

All zebrafish strains were maintained alternating generations of incrosses and outcrosses with wild type. Embryos were obtained by mating adult fish following standard methods and grown at 28.5°C. 1% 1-phenyl-2-thiourea (PTU) (Sigma-Aldrich) was used as an inhibitor of pigmentation from 24hpf onward. The Tg[βactin:HRAS-EGFP] line (termed Tg[CAAX:GFP] herein) expresses GFP in the plasma membrane and was used to label the cell contours^46^. Tg[vglut2a:Lox:DsRed:Lox-GFP] and Tg[gad1b:Lox:DsRed:Lox:GFP] switch lines label glutamatergic and GABAergic neurons, respectively^26^. The Tg[HuC:GFP] line labels differentiated neurons^47^. The Tg[elavl3:jrGECO1b] transgenic line is a reporter of neuronal activity^48^. Tg[neurog1:CreP2A-nlsCerulean] transgenic line was generated using Tol2 transgenesis.

### CRISPR/Cas9-based approach to target proneural genes in F_0_

Four different CRISPR RNAs guides (gRNAs) were used to redundantly target PN genes following^34^. The strategy was to target 4 sites of each PN gene within the bHLH DNA binding domain coding sequence to either generate a frame shift or a premature stop codon either prior to the DNA binding sequence or within the DNA binding sequence, or alternatively a complete excision of the DNA binding sequence (Figure S3b). As controls, we used 4 gRNAs not targeting the zebrafish genome designed by randomization of antibiotic resistance gene sequences. All CRISPR RNAs (crRNAs) were ordered from Integrated DNA Technologies (IDT) (Table S1). One-cell stage embryos were injected with a Cas9/gRNAs mix, containing IDT Alt-R HiFi S.p. Cas9 Nuclease V3 (62mM) and a 4 gRNAs mix made from IDT crRNA and tracrRNA annealing (100mM). Embryos were injected with ∼1nl of 9mM RNP with a 1:1 Ratio of Cas9 and gRNA incubated at 37°C for 5min and chilled on ice before injection^34^. Deletion efficiency was assessed by gDNA PCR amplification of injected embryos (see Table S2 for primers). Disruption of the coding sequence (CDS) was considered complete when no band of the original sequence length was observed, and partial when a faint band of the original sequence length was still visible. Combinations of gRNAs were tested until they led to most of the samples (n=10) displaying complete disruption. For lineage analyses and models we used n=10 embryos, and for calcium imaging n=7 embryos per control condition and n=3-4 embryos per CRISPANT condition.

### Whole mount Hybridization Chain Reaction

Zebrafish embryos at selected stages were fixed overnight in 4% paraformaldehyde (PFA) at 4 °C, dehydrated through a methanol series (25%, 50%, 75%, 100%), and stored at −20 °C until use. After rehydration, embryos were permeabilized with proteinase K (10 μg/ml in PBST; PBS with 0.1% Tween-20) for 10 min at 24hpf or 20 min at 48hpf, followed by post-fixation in 4% PFA for 20 min at room temperature. HCR™ Gold RNA-FISH was performed according to the manufacturer’s instructions (Molecular Instruments, Los Angeles, CA). Custom probe sets targeting *neurog1* (X1), *ptf1a* (X2), *atoh1a* (X7), *Cre* (X7), and *ascl1b* (X3) were used with corresponding amplifiers (X1–488, X2–546, X7–514, X3–647). Hybridization was carried out overnight, followed by overnight incubation with amplifiers. Embryos were imaged within 3 days using a SP8 Leica inverted confocal microscope.

### TUNEL assay

Distribution of apoptotic cells was determined by TdT-mediated dUTP nick-end labeling of the fragmented DNA (TUNEL, Roche). Briefly, whole embryos at 24hpf or 72hpf were fixed in 4% PFA and dehydrated in 100% MetOH. They were permeabilized with Proteinase K at 25 μg/ml and preincubated with TUNEL mixture during 60 min at 37°C according to the manufacturer’s instructions. DAPI (1:500; Molecular Probes) was used to label all nuclei.

### Confocal imaging of whole mount embryos

Zebrafish embryos at the desired time were anesthetized with tricaine and mounted dorsally in 1% low melting point agarose with the hindbrain positioned towards the glass-bottom Petri dish (Mattek). Images were acquired with either a Leica SP8 or Stellaris8 system using 20×glycerol immersion objective, HCX PL APO Lambda blue 20×/0.7 argon laser 30%-80%, DPSS 561 Ready Lasers. z-stacks were recorded with 0.57×0.57×1.04 μm voxel size. The gain was adjusted to each signal to cover the full range of pixel values, minimize signal to noise ratio and compensate inter-individual variability.

### Confocal image processing

Briefly, confocal images were processed using a standardized workflow in which they were 1) deconvoluted using a Richardson-Lucy algorithm and a theoretical PSF, 2) cropped and flipped to be in a consistent orientation and to reduce weight of duplicated images, 3) aligned to a common reference using our rigid alignment software (RAD), 4) segmented at the single cell level using Cellpose^35,49^, 5) masks were intersected to identify cells that strictly expressed Cre prior to the neurotransmitter and cell count was obtained, and 6) masks of 10 embryos were averaged to generate a population spatial distribution 3D model.

### Light Sheet Fluorescence Microscopy

For developmental recordings double transgenic Tg[neurog1:Cre;vglut2:switch] and Tg[neurog1:Cre;gad1b:switch] embryos were anesthetized with tricaine and mounted in 0.35% LMP-agarose within a V-shape chamber filled with fish water. Imaging was performed every 5 min for 24h at 28.5°C on a Viventis microscope using 16× objective. 445nm, 488nm and 560nm channels were simultaneously recorded for 3 embryos every 5 min.

For calcium imaging, Tg[elavl3:jrGeco1b] embryos were screened for the fluorescent transgenic indicators, immobilized by immersion in E3 water containing alpha-bungarotoxin (1mg/mL) for 2min and rinsed in E3 water for 20min until unresponsive to touch^50,51^. Embryos were then embedded in 2% low melting point agarose (Type VII, Si gma-Aldrich) prepared in filtered fish water and encased within a custom-designed glass capillary (2 mm outer diameter, 20 mm length; Hilgenberg GmbH). The agarose encased embryo was then partially extruded out of the capillary to expose the embryo head. The capillary itself was mounted vertically in the imaging specimen chamber of a SiMView light-sheet microscope filled with filtered fish water maintained at 28.5°C, with the dorsal side of the embryo facing the detection objective (Nikon 16x/0.8 NA). A volume of ∼800×400×200μm^3^ (X×Y×Z) was acquired with a Z spacing of 2.2μm at a rate of 2Hz, using a 5ms exposure time per image and an illumination laser wavelength of 561nm with 4mW beam power and 2.5µm beam diameter at the sample location. Embryos were recorded for 30min.

### Calcium Imaging analysis

Each data set was processed with a MATLAB pipeline previously described^52^, and then further processed and integrated with our other datasets as follows. Briefly, recordings were first converted from the binary file format enabling such high-speed data acquisition to conventional .tiff files, a rolling window defined reference of intensity at regular intervals was used for further ΔF/F computations, and finally ΔF/F data sets were filtered using a median filter of kernel 3 to remove noise. Using a Fiji macro, they were cropped to save storage space and simplify downstream processing, as well as standardized their magnification with respect to our confocal datasets. For the resulting recordings, the first frame was then used to perform the alignment, and the transformation matrix was then automatically applied to each frame of the recording within our alignment software (RAD). We then generated a projection over the temporal dimension resulting in a 3D stack representing the most intensely active neurons. This stack was then used to perform a single cell segmentation using Cellpose. Then, individual cell masks were used to trace the fluctuations in intensity across time at those coordinates. Each cell’s track was de-trended with a running baseline F0 that follows the low signal bleaching/drift. Synchrony was measured as the mean absolute Pearson correlation of a cell’s ΔF/F with every other cell of the embryo, whereas coherence of population was measured as the Pearson correlation of a cell’s ΔF/F with every other cell within that group. Event rate was measured as the number of events per minute, and event amplitude the mean peak height. Control metrics where quantified to ensure that measured neuronal activity features differences between LOF of function experiments were not the result of variation of baseline F0 (Figure S6f), detected active neuron per embryos (Figure S6g), total imaged volume (Figure S6h) or lower SNR (Figure S6i).

### Rigid Alignment with Deep Learning (RAD)

To align images from multiple samples and modalities we developed a Python-based interface with integrated neural network and manual 3D rigid registration capabilities. In brief, it leverages the SITK Python library to apply the computed transformation component generated either through a Python based neural network or through manual user input. It offers different possibilities for neural network training, either leveraging manual inputs or a folder of original and pre-aligned images. It is designed to handle the 3D rigid registration of samples batches as well as the drift correction and jitter correction of 3D timeseries. A set of previously aligned images^15^ was used to prepare a training dataset containing about 40 combinations of reference and aligned images. The training dataset was further amplified 5-fold using random transformations (within the range of traditional alignment) allowing to largely extend the range of transformations visible to the neural network during training. The trained model was then used to process each alignment, and each registration was observed and manually adjusted within the UI if deemed necessary before being saved. When performing alignment across modality, a first sample was often aligned manually and then used as reference for other samples of that modality to spare the training of a cross-modality model.

### Statistics and reproducibility

Cell counts were obtained using single cell segmentation mask from Cellpose. The trained Cellpose model and script used to generate the training dataset are available upon request. Statistical analysis was performed in GraphPad Prism 9. For each quantified experiment, the number of replicates/embryos (n) and P values are indicated in the corresponding Supplementary Dataset 1. The experiments reported here were repeated independently at least three times to generate the analyzed dataset. Data distribution was not assumed to be normal; a homoscedasticity test was performed prior to the statistical test selection. A z-score mediated outlier detection across all measured cell counts was used to rule out samples from an original pool containing from 5 to 13 embryos for each condition and transgenic background. Data analyses were performed using automated procedures without consideration of experimental groups. Data analysis was performed using the same parameters across groups.

For the calcium analysis and quantification, the strategy was similar except for the shapely variance attribution. Briefly, for each of the control embryos, we interrogated how well three atlas maps (neurotransmitter, lineage and birthdate) predicted the neuronal activity feature at the same registered position in other embryos. The positional ceiling C was the R² of predicting each cell by the mean of its 50 nearest cells in the other embryos. The R² of all three maps was divided among them by Shapley values^53^, which equate to the general dominance weights and have been applied to cross-validated predictiveness^54^. Plots show each map’s share of the positionally reproducible variance, Shapley share/C, per embryo.

## ACKNOWLEDGEMENTS

We would like to thank Dulce Real for technical assistance and Dr Daniel Feliciano and members of the Espinosa-Medina’s and Pujades’ labs for insights and critical discussions. We thank and Dr Kyle Loh for critical reading of the manuscript. We thank the Janelia Visiting Scientist Program for supporting the collaboration between HHMI Janelia Research Campus and UPF labs, and Dr Misha Arehns’ lab for sharing the Tg[elavl3:jrGECO1b] transgenic line. The authors thank the Advanced Light Microscopy Unit at the CRG where confocal imaging was performed, the Scientific Computing Core Facility (MELIS-UPF), and the Aquatic Core Facilities at both UPF-PRBB and Janelia. This work was funded by grants PID2021-123261NB-I00 and PID2024-160356NB-I00 from Ministerio de Ciencia y Universidades (MICIU), Agencia Estatal de Investigación (AEI, DOI: 10.13039/501100011033) and Fondo Europeo de Desarrollo Regional (FEDER) to CP. The Department of Medicine and Life Sciences (UPF) is a Unidad de Excelencia María de Maeztu (CEX2018-000792-M) funded by the AEI. LM was a recipient of a EUTOPIA-SIF postdoctoral fellowship. GOÁ is a recipient of a post-doctoral fellowship from “La Caixa” Foundation (ID 100010434; code LCF/BQ/PR23/11980036). CP is a recipient of ICREA Academia award (Generalitat de Catalunya).

## AUTHOR CONTRIBUTIONS

Conceptualization: MB, IEM and CP; Methodology: MB, LM, GOA, WCL, PK; Investigation and formal analysis: MB; Validation and analysis: MB and CP; Resources: PK; Writing-original draft: MB, IEM and CP; Visualization: MB; Supervision: CP; Project administration: CP; Funding acquisition: IEM and CP.

## DATA, CODE AND MATERIALS AVAILABILITY

All relevant data can be found within the article and its supplementary information. Code is freely available in https://github.com/cristinapujades/Blanc_et_al_2025. For more information or any specific tool, you should contact us.

## SUPPLEMENTARY FIGURES’ LEGENDS

**Figure S1:**
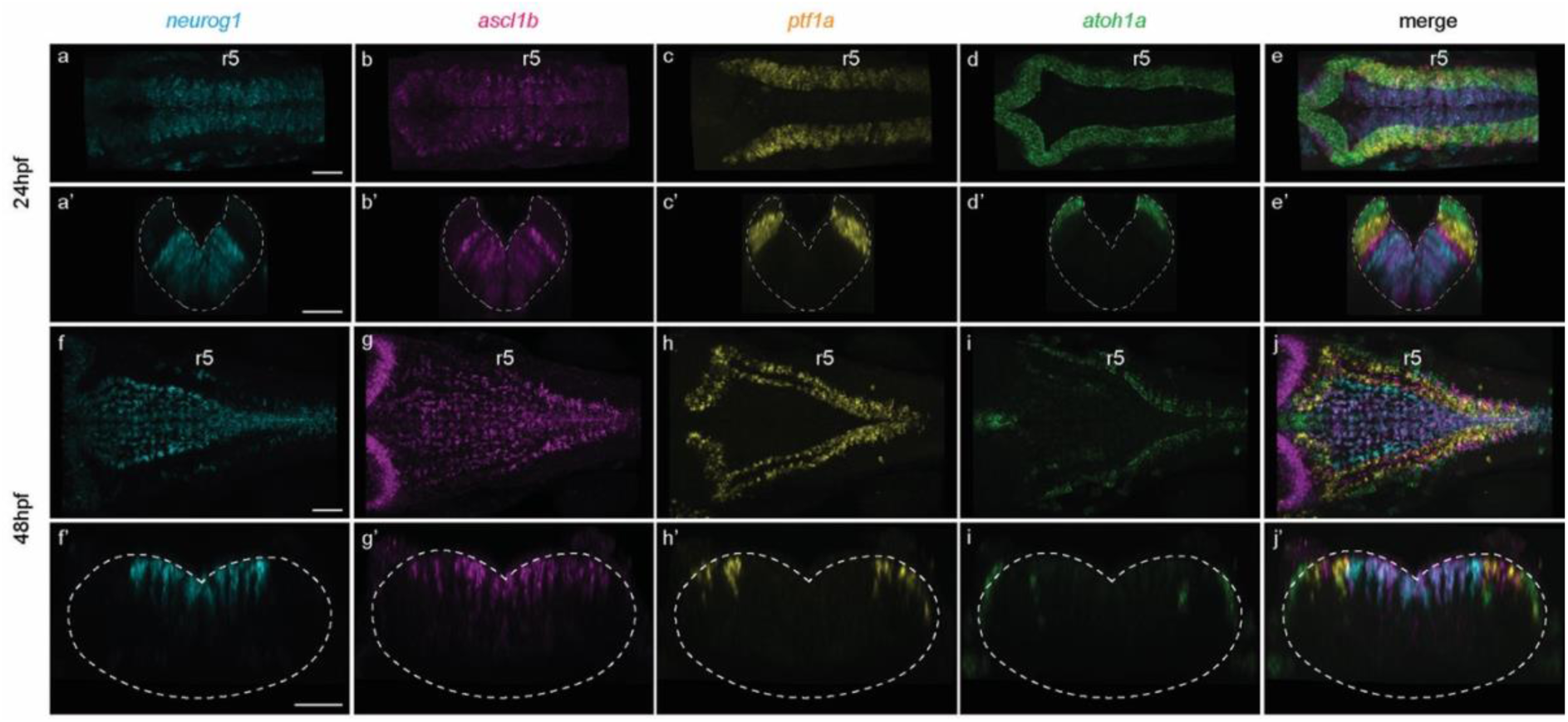
Proneural genes are differentially expressed along the DV and ML axis within the hindbrain. a–j, a’–j’, Multiplex HCR of *neurog1*, *ascl1b*, *ptf1a*, *atoh1a* proneural genes displaying single (a–d, f–i) and merged (e, j) channels at 24hpf (a–e, a’–e’) and 48hpf (f–j, f’–j’), in dorsal (a–j) and transverse (a’–j’) views along r3. Note that proneural genes are expressed in the progenitor domain and their expression is displaced upon the growth of the below neuronal differentiation domain. hpf, hours post-fertilization; r, rhombomere.

**Figure S2:**
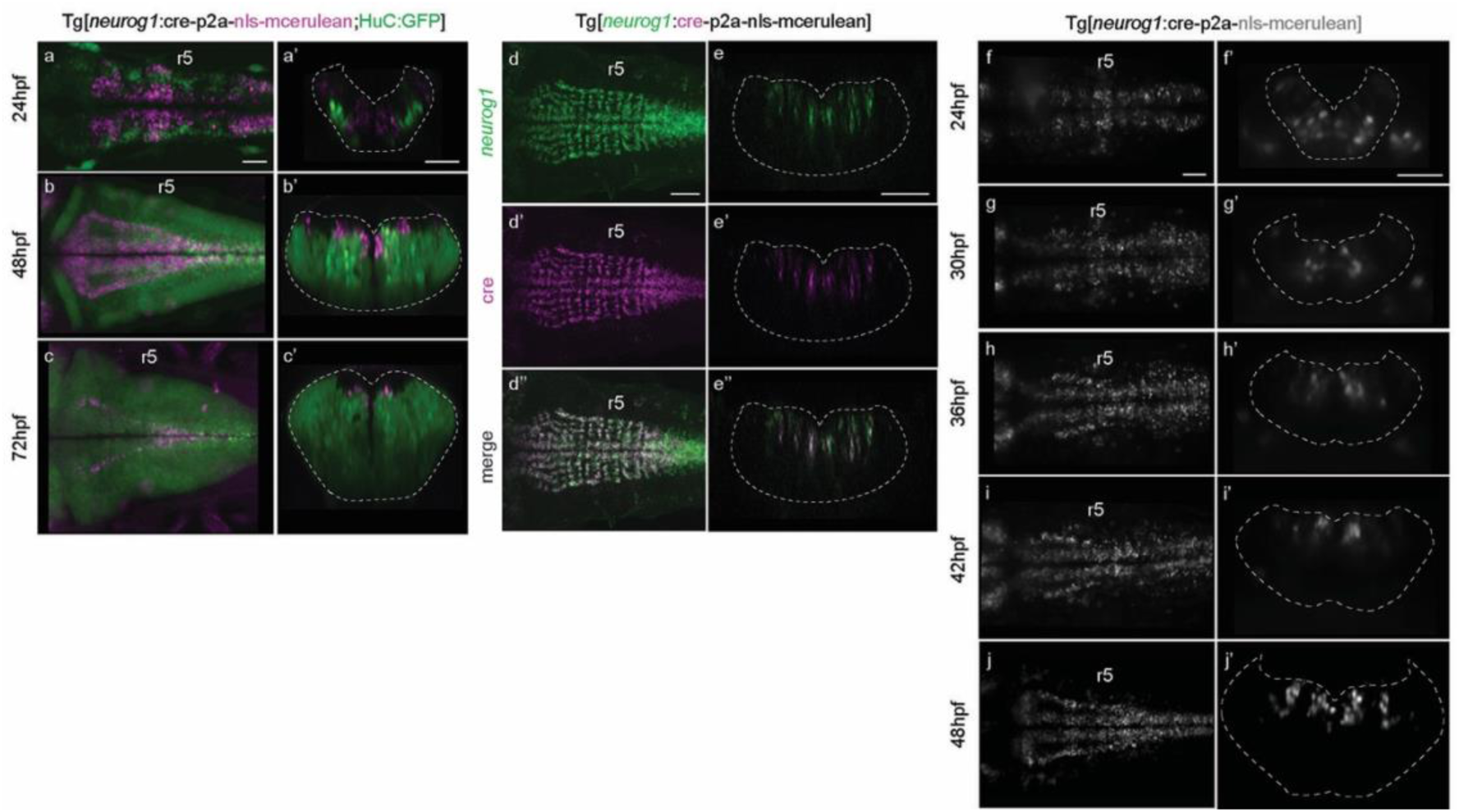
Characterization of the Tg[neurog1:Cre-P2A-nls-Cerulean] fish line. a–c, a’–c’, Double transgenic Tg[neurog1:Cre-P2A-nls-Cerulean;HuC:GFP] embryos at the indicated developmental stages. Dorsal maximal intensity projection (MIP) (a–c) and transverse view through rhombomere 5 (a’–c’). Note that neurog1-Cerulean expression in magenta is restricted to the progenitor domain. d–d’’, e–e’’, HCR for *neurog1* and *Cre* in Tg[neurog1:Cre-P2A-nls-Cerulean] embryos at 40hpf. Note that *Cre*-positive cells overlap with *neurog1*-cells. f–j, f’–j’, Stills from a time-lapse video showing the expression of Cerulean protein recapitulates the dynamics of neurog1 expression^27^. Note the stability of Cerulean protein, which is maintained even in neurons deriving from neurog1-cells. a–c, d–d’’, f–j, Dorsal views with anterior to the left. a’–c’, e–e’’, f’–j’, Transverse views through rhombomere 5. r, rhombomere; hpf, hours post-fertilization.

**Figure S3:**
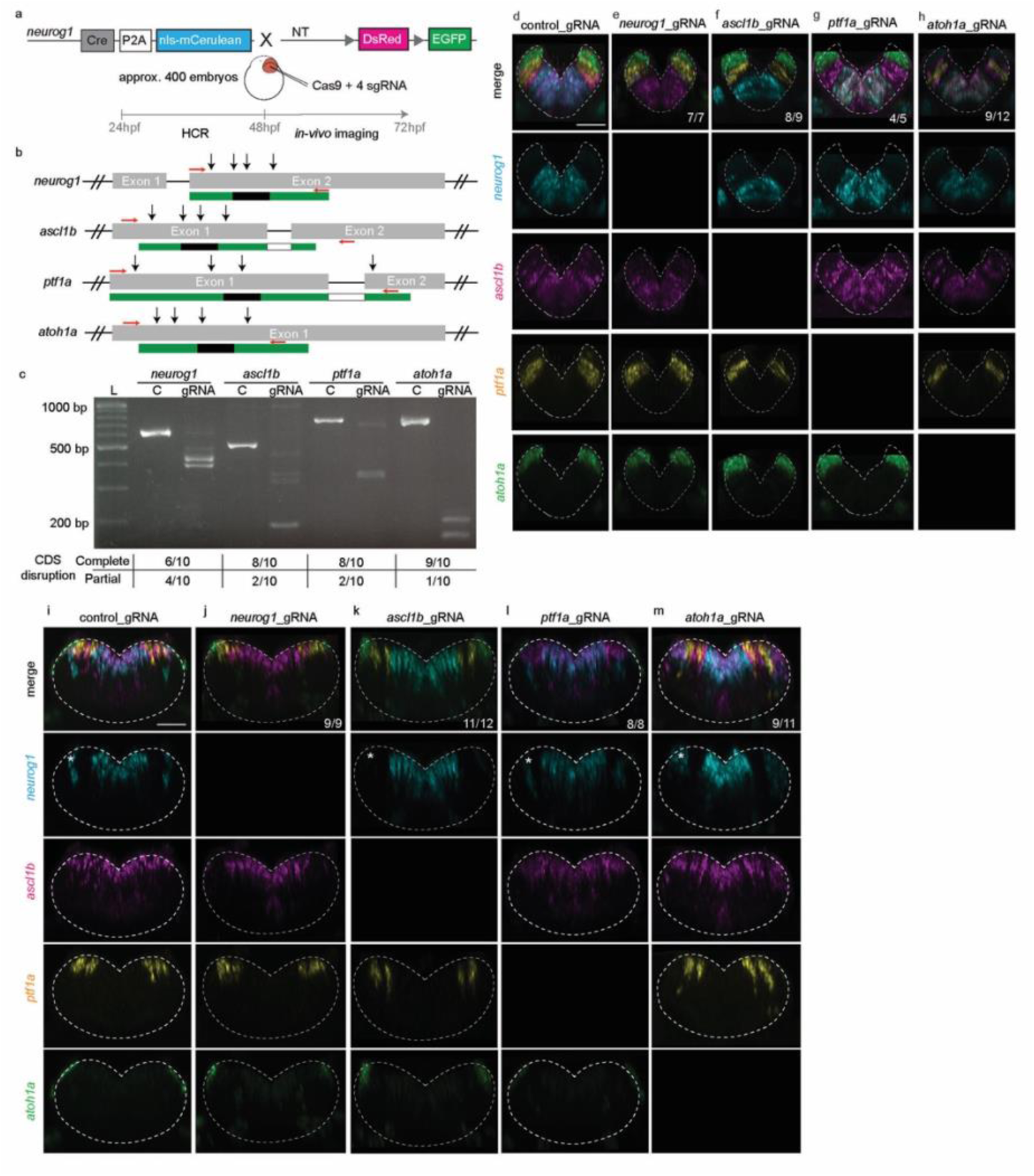
CRISPR-based loss-of-function approach. a, Scheme depicting the CRISPR/Cas9-based approach that produces loss-of-function (LOF) phenotypes in F_0_. Tg[neurog1:Cre:vglut2a:switch] and Tg[neurog1:Cre:gad1b:switch] embryos were injected with control or 4 different gRNAs targeting a single proneural gene (PN_gRNAs). They were let to develop until 24hpf and 48hpf for HCR proneural gene expression analysis, and 72hpf for cell lineage studies (Figure 2). b, Strategy for the design of gRNAs targeting proneural genes. Black arrows indicate the targeting position of the 4 gRNAs within the corresponding proneural gene. Red arrows indicate the position of the PCR primers used for genotyping (Table S1). c, Genotyping of PN_gRNAs injected embryos. Genomic DNA from single embryos injected with control (C) or PN_gRNAs (gRNA) was extracted and used to assess disruption in the coding sequence (CDS) by PCR using specific primers (red arrows). Agarose gel displaying the amplified bands. The number of embryos displaying complete or partial disruption of the amplified coding sequence (CDS) after PN_gRNAs injection is indicated in the below table. Efficiency of the gene disruption was assessed in single embryos at F_0_. d–h, i–m, Multiplex HCR of *neurog1*, *ascl1b*, *ptf1a* and *atoh1a* proneural genes at 24hpf and 48hpf, respectively, upon control (d, i) or PN_gRNAs injection (e–h, j–m). Transverse views through rhombomere 5 displaying the indicated single channel or the merge of them. White asterisk in the *neurog1*-profiled embryos (i–m) indicates the expression of *neurog1*-in the most lateral domain of expression. Note that this domain disappears upon *ascl1b* loss-of-function.

**Figure S4:**
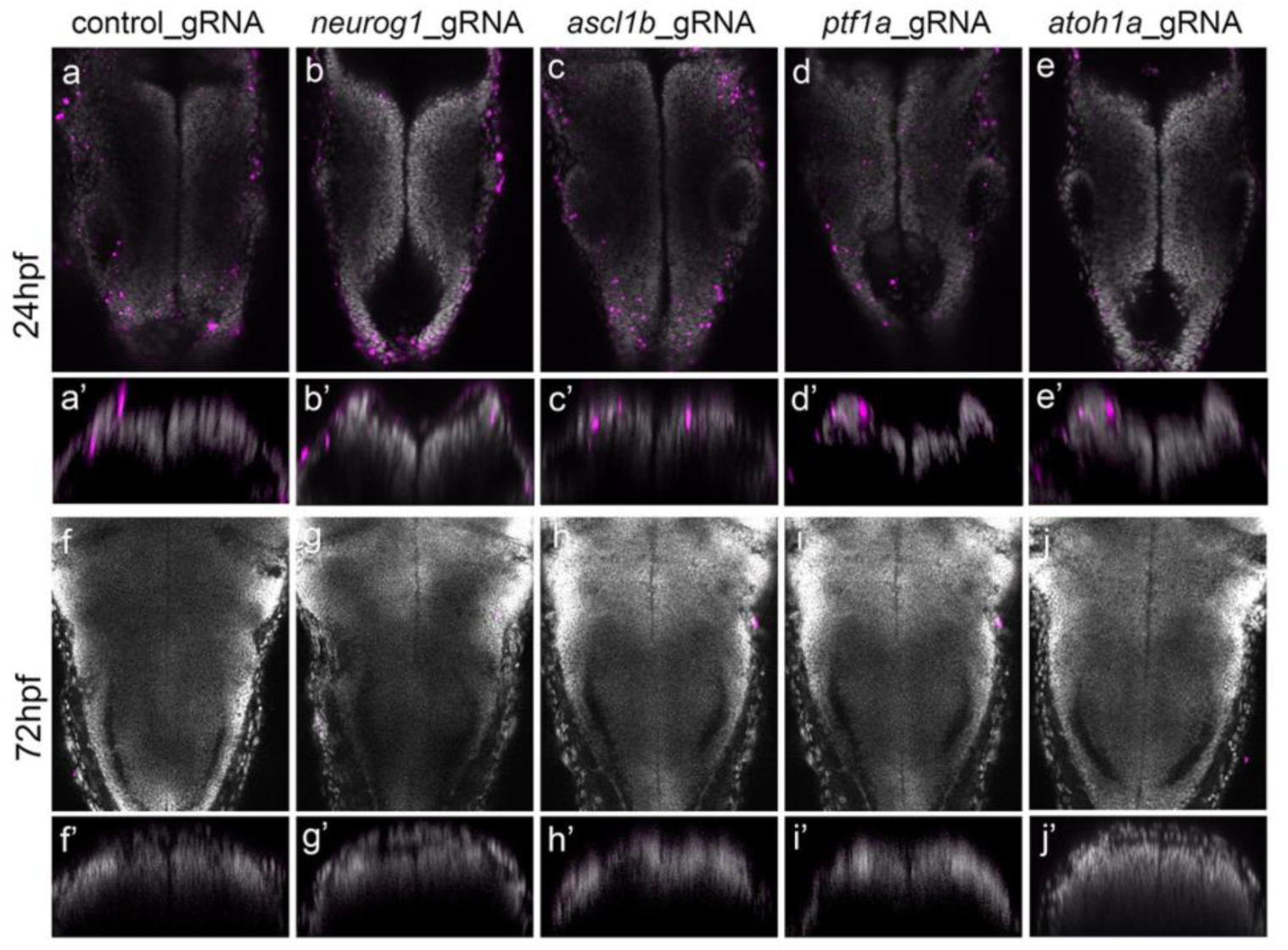
Cell apoptosis analysis in CRISPANT embryos. a–j, Tg[neurog1:Cre:vglut2a:switch] embryos were injected with control or 4 different gRNAs targeting a single proneural gene (PN_gRNAs) and let to develop until 24hpf (a–e, a’–e’) and 72hpf (f–j, f’–j’) for TUNEL assay (magenta cells) and DAPI staining (grey). a–j, Dorsal views with anterior to the top. a’–j’, Transverse views.

**Figure S5:**
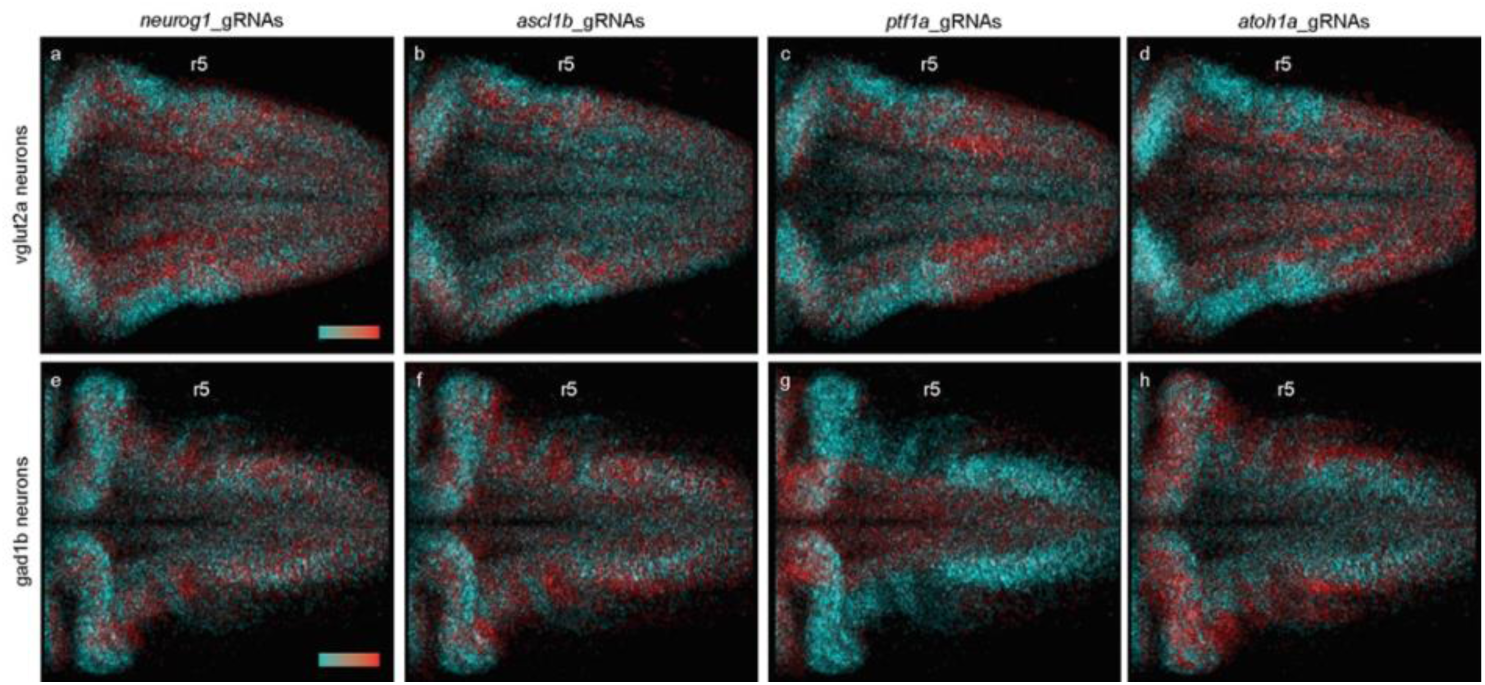
Differential requirement of progenitor pools for glutamatergic and GABAergic lineages. a–h, 3D-maps displaying the progenitor origin requirement (see cyan in hue legend) and compensation (see red in hue legend) for glutamatergic (a–d) and GABAergic (e–h) neurons. Note that the main progenitor’s requirements for glutamatergic neurons are *ascl1b*-(medial) and *atoh1a*-(lateral), whereas to the GABAergic neurons are *ptf1a*-progenitors. All images display dorsal MIP with anterior to the left. r, rhombomere.

**Figure S6:**
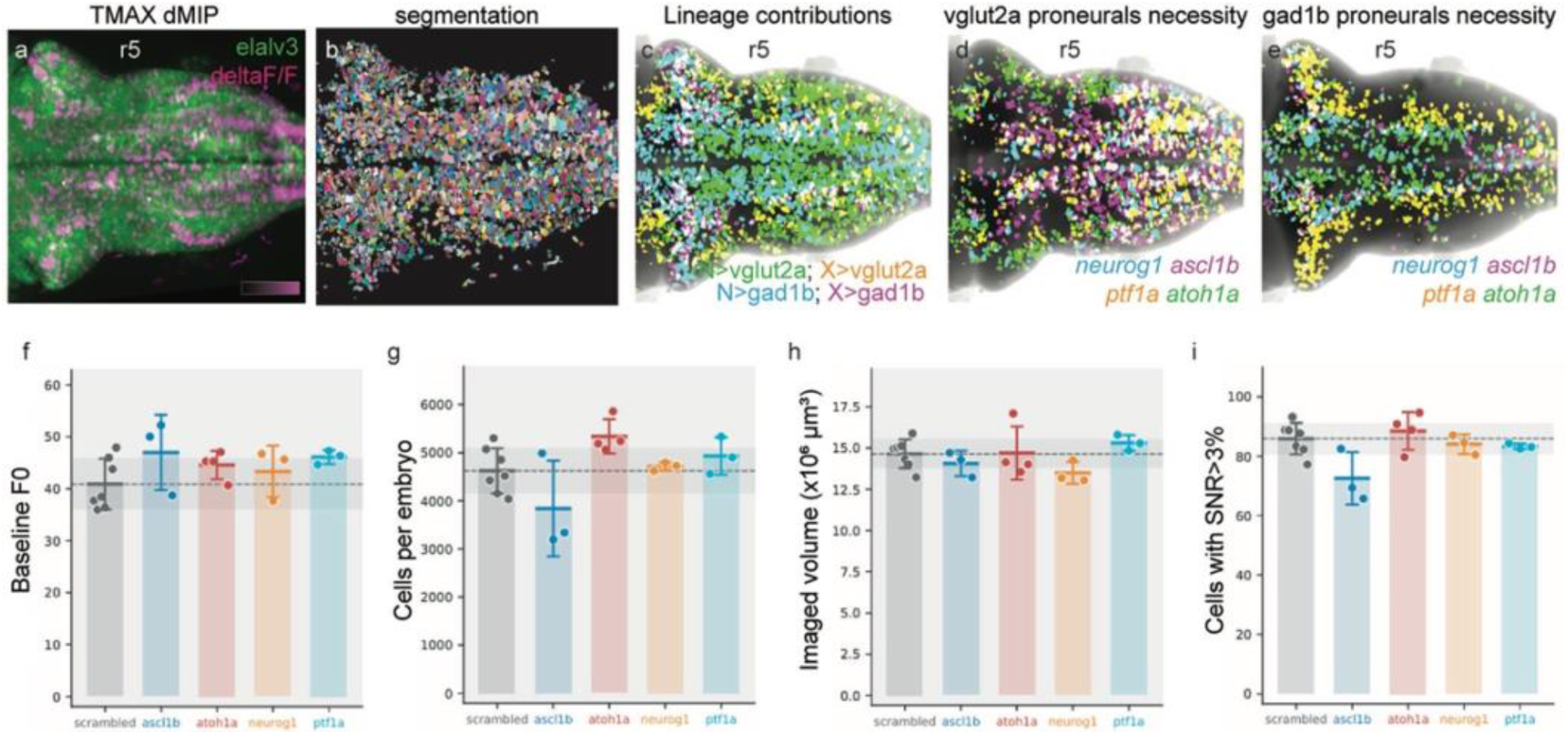
Mapping lineage origin and neuronal activity features to a common hindbrain 3D reference map. a, Dorsal view of the hindbrain of a Tg[elavl3:jrGECO1b] embryo upon calcium-recording at 72hpf (Video 6). b, Dorsal view displaying all segmented active neurons. c, 3D-map displaying the contribution of *neurog1*-progenitors (green and cyan cells) and non *neurog1*-progenitors (yellow and magenta cells) to glutamatergic (green and yellow) and GABAergic (cyan and magenta) lineages. d–e, 3D-maps of the proneural gene requirements for glutamatergic (d) and GABAergic (e) lineages. f–i, Histogram displaying the average baseline F0 values, number of cells, imaged volume and cells with signal to noise ratio superior to 3% among all samples and conditions.

## VIDEOS

**Video 1: *neurog1*-population is the main contributor to hindbrain glutamatergic neurons at very early embryonic stages.**

Tg[neurog1:Cre:vglut:switch] embryo was imaged from 24hpf until 48hpf using LSFM. Note that *neurog1*-progenitors express Cerulean and when they differentiate into glutamatergic neurons, they start expressing GFP (green), whereas the rest of glutamatergic neurons express DsRed (magenta). Dorsal view with anterior to the left.

**Video 2: *neurog1*-population and other progenitor pools contribute to glutamatergic neurons at later embryonic stages.**

Tg[neurog1:Cre:vglut:switch] embryo was imaged from 48hpf until 72hpf using LSFM. Note that *neurog1*-progenitors express Cerulean and when they differentiate into glutamatergic neurons, they start expressing GFP (green), whereas the rest of glutamatergic neurons express DsRed (magenta). Dorsal view with anterior to the left.

**Video 3: *neurog1*-population produces GABAergic neurons at early embryonic stages.**

Tg[neurog1:Cre:gad1b:switch] embryo was imaged from 24hpf until 48hpf using LSFM. Note that *neurog1*-progenitors express Cerulean and when they differentiate into GABAergic neurons, they start expressing GFP (green), whereas the rest of GABAergic neurons express DsRed (magenta). Dorsal view with anterior to the left.

**Video 4: *neurog1*-population and other progenitor pools produce GABAergic neurons at later embryonic stages.**

Tg[neurog1:Cre:gad1b:switch] embryo was imaged from 48hpf until 72hpf using LSFM. Note that *non neurog1*-progenitors (magenta cells) take the relay of neurog1-progenitors in the production of GABAergic neurons. Dorsal view with anterior to the left.

**Video 5: GABAergic neurons are generated independently of *ptf1a*.**

Tg[neurog1:Cre:gad1b:switch] embryo injected with 4 different gRNAs for *ptf1a* was imaged from 48hpf until 72hpf using LSFM. Note that *neurog1*-cells (cerulean) give rise to GABAergic neurons (green) in the absence of *ptf1a*. Dorsal view with anterior to the left.

**Video 6: Neuronal activity pattern in the embryonic hindbrain at 72hpf.**

Calcium recordings in Tg[elavl3:jrGECO1b] embryos at 72hpf after control_gRNAs injection. Dorsal view with anterior to the left.

**Video 7: Neuronal activity pattern in the hindbrain upon disruption of *neurog1*-progenitors.**

Calcium recordings in Tg[elavl3:jrGECO1b] embryo at 72hpf after *neurog1*_gRNAs injection. Note how the Ca^++^-activity pattern changes upon disruption of *neurog1*-progenitors. Dorsal view with anterior to the left.

**Video 8: Neuronal activity pattern in the hindbrain upon disruption of *ascl1b*-progenitors.**

Calcium recordings in Tg[elavl3:jrGECO1b] embryo at 72hpf after *ascl1b*_gRNAs injection. Note how the calcium-activity pattern changes upon disruption of *ascl1b*-progenitors. Dorsal view with anterior to the left.

**Video 9: Neuronal activity pattern in the hindbrain upon disruption of *ptf1a*-progenitors.**

Calcium recordings in Tg[elavl3:jrGECO1b] embryo at 72hpf after *ptf1a*_gRNAs injection. Note how the calcium-activity pattern changes upon disruption of *ptf1a*-progenitors. Dorsal view with anterior to the left.

**Video 10: Neuronal activity pattern in the hindbrain upon disruption of *atoh1a*-progenitors.**

Calcium recordings in Tg[elavl3:jrGECO1b] embryo at 72hpf after *atoh1a*_gRNAs injection. Note how the calcium-activity pattern changes upon disruption of *atoh1a*-progenitors. Dorsal view with anterior to the left.

---

**Table S1.**
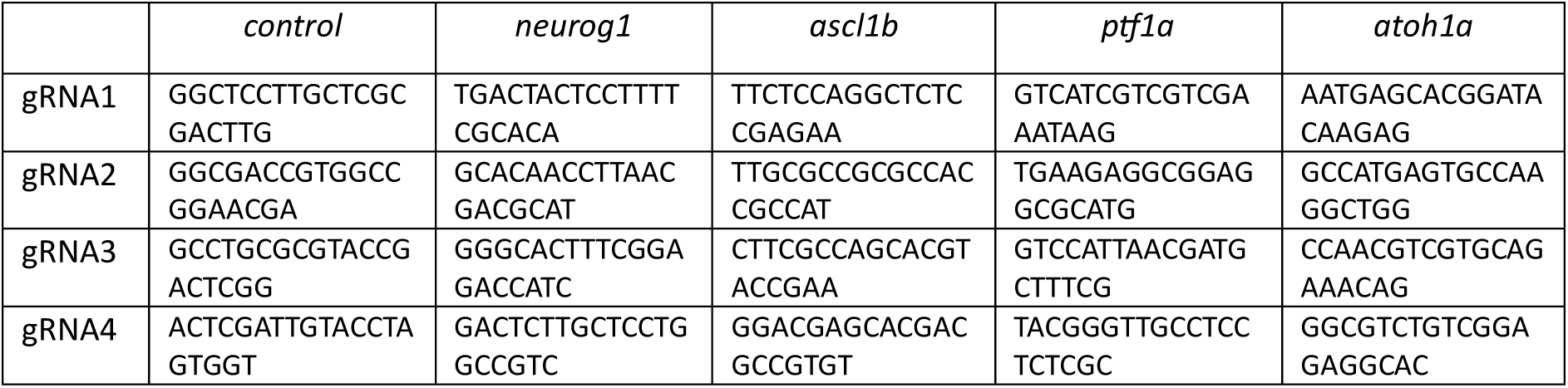
Sequences of CRISPR RNAs (crRNAs) used to target proneural genes (5’ – 3’)

**Table S2.**
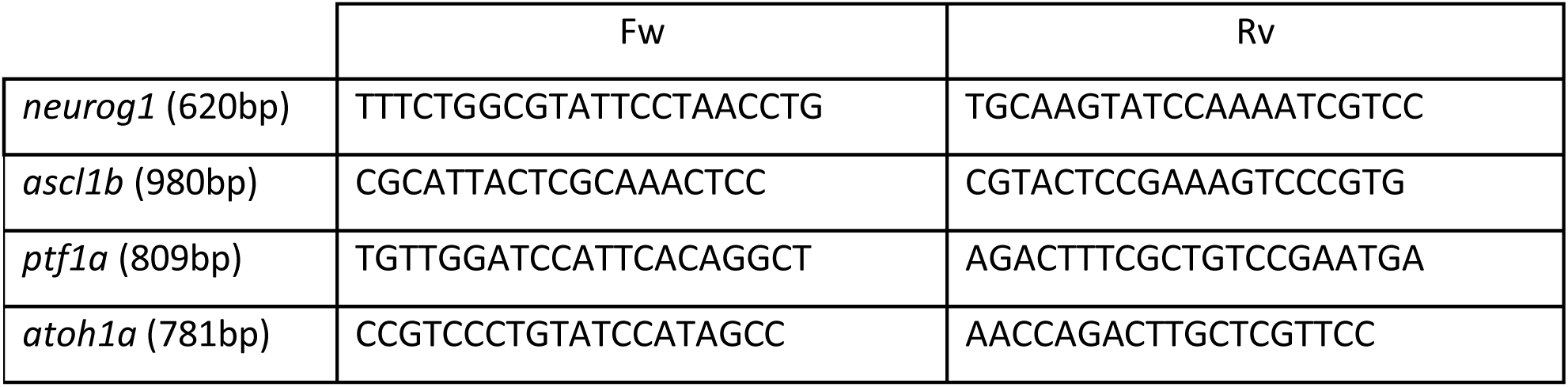
Sequence of oligonucleotides used for proneural genes PCR amplification of gDNA from injected embryos to assess deletion efficiency (5’ – 3’)

